# Establishing a Living Biobank of Patient-Derived Organoids of Intraductal Papillary Mucinous Neoplasms of the Pancreas

**DOI:** 10.1101/2020.09.11.283168

**Authors:** Francisca Beato, Dayana Reverón, Kaleena B. Dezsi, Antonio Ortiz, Joseph O. Johnson, Dung-Tsa Chen, Karla Ali, Sean J. Yoder, Daniel Jeong, Mokenge Malafa, Pamela Hodul, Kun Jiang, Barbara A. Centeno, Mahmoud A. Abdalah, Jodi A. Balasi, Alexandra F. Tassielli, Bhaswati Sarcar, Jamie K. Teer, Gina M. DeNicola, Jennifer B. Permuth, Jason B. Fleming

## Abstract

Pancreatic cancer (PaCa) is the third leading cause of cancer-related deaths in the United States. There is an unmet need to develop strategies to detect PaCa at an early, operable stage and prevent its progression. Intraductal papillary mucinous neoplasms (IPMNs) are cystic PaCa precursors that comprise nearly 50% of pancreatic cysts detected incidentally via cross-sectional imaging. Since IPMNs can progress from low- and moderate-grade dysplasia to high-grade dysplasia and invasion, the study of these lesions offers a prime opportunity to develop early detection and prevention strategies. Organoids are an ideal preclinical platform to study IPMNs, and the objective of the current investigation was to establish a living biobank of patient-derived organoids (PDO) from IPMNs. IPMN tumors and adjacent normal pancreatic tissues were successfully harvested from 15 patients with IPMNs undergoing pancreatic surgical resection at Moffitt Cancer Center & Research Institute (Tampa, FL) between May of 2017 and March of 2019. Organoid cultures were also generated from cryopreserved tissues. Organoid count and size were determined over time by both Image-Pro Premier 3D Version 9.1 digital platform and Matlab application of a Circular Hough Transform algorithm, and histologic and genomic characterization of a subset of the organoids was performed using immunohistochemistry and targeted sequencing, respectively. The success rates for organoid generation from IPMN tumor and adjacent normal pancreatic tissues were 81% and 87%, respectively. IPMN organoids derived from different epithelial subtypes showed different morphologies *in vitro*, and organoids recapitulated histologic and genomic characteristics of the parental IPMN tumor. In summary, this pre-clinical model has the potential to provide new opportunities to unveil mechanisms of IPMN progression to invasion and to shed insight into novel biomarkers for early detection and targets for chemoprevention.

## Introduction

Pancreatic cancer (PaCa) poses a significant mortality burden globally. It is currently the seventh leading cause of cancer-related deaths worldwide^1^ and the third leading cause in the United States (US)^2^. Approximately 57,600 patients will be diagnosed with pancreatic cancer this year in the US and an estimated 47,050 patients will die from this malignancy^3^. Despite advances in its diagnosis and treatment, it holds the lowest five-year relative survival rate of all leading cancers, at 9-10%^3^. PaCa incidence and mortality rates are increasing^4-8^, and it is predicted to become the second leading cause of cancer mortality around 2020, surpassing colorectal, prostate and breast cancers^9^. There is an unmet need to develop strategies to detect PaCa at an early, operable stage and prevent its progression.

Intraductal papillary mucinous neoplasms (IPMNs) are cystic PaCa precursors that comprise nearly 50% of pancreatic cysts detected incidentally via computed tomography (CT) scans and magnetic resonance imaging (MRI)^10, 11^. Unfortunately, once detected, noninvasive strategies to accurately distinguish benign IPMNs that can undergo surveillance from malignant IPMNs that warrant surgical resection are lacking, posing a clinical conundrum. IPMNs can present in the main pancreatic duct (MD-IPMN), side branch ducts (BD-IPMN), or both (mixed-IPMN). Based on cytoarchitectural features and mucin protein profiles assessed pathologically after tissue has been resected, IPMNs are classified into four histopathological types: gastric, intestinal, pancreatobiliary, and oncocytic, with the pancreatobiliary subtype having the poorest prognosis^12, 13^.

Up to 70% of resected MD -IPMNs harbor high-grade dysplasia or invasive disease^14, 15^ and have a risk of recurrence up to 65% if invasive disease is present^16^. Of patients diagnosed with well- and poorly-differentiated unresectable IPMNs, clinical surveillance studies show malignant progression within 10 years in up to 8% and 25% of cases, respectively^15, 16,17^. Taken together, because IPMNs can progress to invasive, fatal malignancy, the study of these lesions offers a prime opportunity to develop early detection and prevention strategies.

The organoid culture approach provides a potential framework for preclinical and clinical PaCa research focused on early detection and prevention. This resourceful three-dimensional (3D) culture model recapitulates the structural features, mutational spectrum, tumor heterogeneity and therapeutic sensitivity profiles of primary pancreatic tumors^18-21^. Although patient-derived organoid (PDO) models exist for microscopic pancreatic cancer precursors known as pancreatic intraepithelial neoplasms (PanINs)^22^, our team sought to develop one of the first sets of IPMN PDO models paired with normal tissue. While we were in the process of resubmitting this manuscript, another team^23^ reported on a living IPMN biobank where they generated organoids from 7 normal pancreatic ducts and 10 unpaired IPMN tumor samples.and performed molecular characterization. The aims of the current study were to: (1) establish a human organoid model of paired pre-malignant and normal tissue cultures from resected IPMN specimens; (2) demonstrate the passage and cryopreservation of organoid cultures as part of a living biobank infrastructure; (3) generate pre-invasive organoid cultures from cryopreserved resected pancreatic tissue; (4) establish a reliable protocol for imaging and counting of the organoids via a digital platform; and (5) characterize the primary tumors and organoid cultures by genomic analysis.

## Materials and Methods

### Patient recruitment and tissue collection

Patients diagnosed with resectable pancreatic lesions between May of 2017 and March of 2019 at the Moffitt Cancer Center & Research Institute (Tampa, Florida) were recruited to participate in two complementary Institutional Review Board-approved studies known as the Total Cancer Care Protocol (TCCP) and the Florida Pancreas Collaborative (FPC) study^24^. None of these cases received treatment with chemotherapy or radiation prior to surgery. Tumor and adjacent normal pancreatic tissues were obtained from the surgical suite, harvested, and reviewed by a pancreatic pathologist (BAC or KJ), to verify the diagnosis and to identify tumor and adjacent normal non-neoplastic pancreas tissue. By histology and immunohistochemistry techniques, the epithelial subtype of the IPMN was determined^12, 13^. Both collection and transport of the tissue samples were facilitated by the Moffitt Tissue Core Acquisition Team. The laboratory specialist received the pair of tumor and normal samples within approximately 20 minutes of the operative procedure in order to establish tumor and normal organoid cultures as described below. The workflow from tissue collection to organoid culture characterization is shown in Figure 1.

**Figure 1.**
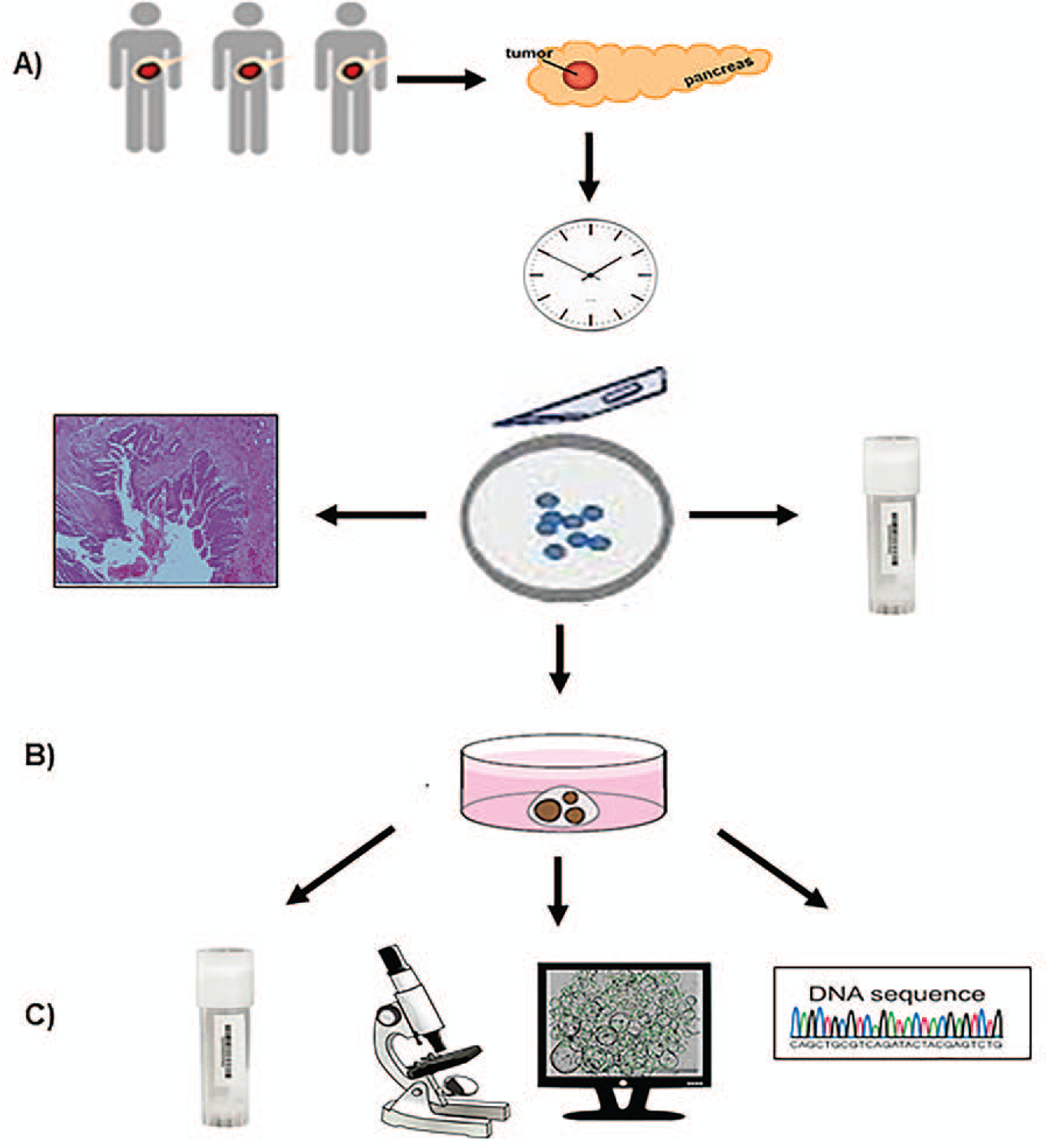
Workflow of patient-derived IPMN organoid culture methodology. A) Resected IPMN tumor and adjacent normal tissues obtained from patients were transported to the processing lab within 20 minutes of resection; and a portion of each type of tissue was cryopreserved. Primary tissue was analyzed by histomorphology and immunohistochemistry analysis. Both IPMN tumor and adjacent normal tissue specimens were minced into small pieces and subsequently subjected to organoid culture. B) 3D-organoid cultures from both tumor and normal tissues were established. C) Organoids were cryopreserved and culture was imaged via bright-field microscopy, automated counting was conducted via the Circular Hough Transform-based algorithm, and characterized by genomic DNA fingerprinting and sequencing.

### Organoid Culture Media

Wash Media consisted of Advanced DMEM/F12 medium supplemented with HEPES [1x, Invitrogen], Glutamine [1x, VWR] or Glutamax [1x, Invitrogen] and Primocin [1x, 50 mg/mL, InvivoGen].

Digestion Media consisted of wash media supplemented with collagenase type IV from Clostridium histolyticum [5 mg/mL, Gibco] and Dispase [1 mg/mL, Sigma].

Complete Growth Media consisted of 50% wash media without FBS, 50% conditioned media [Wnt3a/R-Spondin/Noggin, L-WRN ATCC] supplemented with B27 [1x, Invitrogen], Nicotinamide [10 mM], N-acetyl-L-cysteine [1.25 mM, Sigma], Primocin [100 µg/mL], mNoggin [100 ng/mL], human epidermal growth factor [hEGF, 50 ng/mL, Peprotech], human fibroblast growth factor 10 [100 ng/mL, Peprotech], Gastrin [10 nM, Sigma], A 83-01 [500 nM, Tocris], PGE2 [1 µM, R&D Systems] and Y-27632 [10.5 µM, Sigma].

Cell Recovery Solution (Corning), Recovery Cell culture Freezing Medium (Thermo Fisher).

### Generation and expansion of patient-derived organoids from fresh resected specimens

IPMN tumor and normal pancreatic organoids were generated and expanded by modification of previously-described protocols of PaCa organoids^22, 25-27^. Following resection and receipt of tissues by the lab specialist, tissue samples were placed into separate 50 mL Falcon tubes containing wash media then transferred to a 10 cm dish. A tissue size of 0.25-4 cm^2^ was ideal to move forward with processing. Both tissue types were minced into 1 mm^3^ fragments. Fragments were transferred to a 15 mL tube into which 6 mL of modified digestion media was added. The tube was incubated for 30-45 minutes in a 37°C shaker with agitation at 500 rpm to allow tissue to dissociate into single cells. Fragments were allowed to settle under normal gravity for 2 min and the supernatant was transferred to a clean Falcon tube. An additional 3 mL of wash media containing 10% FBS was added and the tube was centrifuged at 1300 rpm for 5 min. The pellet was washed one more time as above. Cell pellets were then re-suspended in 300 μL of ice-cold growth factor reduced Matrigel. 50 µL domes were loaded onto 6 wells of a pre-warmed 24-well culture plate and allowed to solidify on an incubator for 15 min at 37°C, 5% CO_2_. Next, 500 µL of pre-warmed complete growth media was added per well and incubated at 37°C, 5% CO_2_. The growth and quantity of the organoid cultures were monitored for up to 14 days. Within that period, fresh growth media was added every two days.

At the conclusion of the two-week-incubation, expansion of the organoid cultures was conducted by harvesting from one confluent well. To break apart the Matrigel, 1 mL of cold cell recovery solution or splitting media was added to the well and pipetted up and down. The suspension was incubated in a 15 mL Falcon tube on ice for 30 min, inverted every 10 min, and then centrifuged at 1300 rpm for 6 min at 4°C. The pellet was re-suspended in wash media with 0.1% BSA and washed as above, then re-suspended in Matrigel and plated among 3 to 5 wells of a 24-well plate depending upon organoid size. For example, from a confluent well, smaller organoids (5-50 px.) would be plated into 3 wells, while larger organoids (50-200 px.) would be plated into 5 wells. The culture was allowed to solidify and incubated as described above. The organoid cultures were passaged up to four times.

### Generation and expansion of patient-derived organoids from cryopreserved specimens

Organoids were also derived from cryopreserved specimens with the ultimate goal of determining the viability of pre-malignant organoids cultured from frozen specimens. To accomplish this, following resection, tissue was transferred to a cryovial preloaded with 1 mL of CryoStor CS10 freezing solution (Stem CT) and placed on ice for 30 min, then stored in a Mr. Frosty freezing container at -80 °C overnight prior to liquid nitrogen storage. Tissue was thawed in a 37°C water bath until 50% thawed and transferred to a 100 mm petri dish containing wash media. Tissue was minced and digested as above.

### Freezing and reviving of patient-derived organoids

Organoids from fresh and frozen tissue were harvested using cell recovery solution (1 mL per well), incubated on ice for 30 min, inverted every 10 min and spun at 1300 rpm for 6 min at 4°C. The pellet was re-suspended in wash media with 1% BSA, washed as previously described, and then frozen in 500 µL of cell recovery freezing media. To revive the organoids, the frozen vial was placed in a 37°C water bath until 50% was thawed. The vial was then rinsed with wash media containing 1% BSA and 0.1% Y-27632 Rho kinase inhibitor to help the cells cope with freeze/thaw stress. An additional 3 mL of media were added and the tube was centrifuged at 1300 rpm for 6 min. The resulting pellet was re-suspended in cold Matrigel as previously described. In most cases, the organoids began appearing in culture by three days-post thaw.

### Imaging & digital quantification of patient-derived organoids in vitro

Organoid culture images were acquired on a Zeiss Axio inverted Fluorescence Microscope system using the Zen 2.3 pro analysis and an EVOS FL Auto Imaging System with an Olympus PlanApo N 1.25x/0.04 objective. Z-stacks were obtained on the EVOS FL through the entire volume of the Matrigel in order to capture most of the organoids. Following image acquisition on the EVOS, images were preprocessed and flattened to reduce shadowing effect using Image-Pro Premier Version 9.1. In order to detect the overlapped circular objects, the images were first preprocessed to filter noise, enhance contrast, and enhance the edges. Then, using Matlab a Circular Hough Transform (CHT) based algorithm was used to find the overlapped circular objects in each image. Resulting counts were then plotted and analyzed with the assistance of the Moffitt Analytic Microscopy Core Facility.

### Genomic characterization of patient-derived organoids

#### DNA isolation and quality control

DNA was isolated from both fresh and frozen organoid cultures utilizing the DNeasy Blood & Tissue Kit from Qiagen (Cat. No. 69504). Once DNA was isolated from the organoid cultures, the concentration, volume, and DNA integrity number (DIN) DIN score were determined using Qubit fluorometric quantitation and the Agilent TapeStation Genomic DNA ScreenTape Assay.

#### Preparation of fresh organoids for DNA isolation

To prepare the organoids from culture, the media from one to two confluent wells was aspirated. Cell recovery medium was added (1 mL per well) and pipetted up and down until the Matrigel broke down in solution. The mixture was transferred to a 2 mL tube and incubated on ice for 45–60 min. The mixture was then centrifuged at 6000 RCF for 3 min at 4°C and supernatant was removed followed by re-suspension in 200 µL ice-cold PBS with 0.1% BSA for an initial wash. The solution was triturated vigorously up to 20 times with a P200 pipette. PBS with 0.1% BSA (1 mL) was added to the solution and the tube was centrifuged at 6000 RCF for 3 min at 4°C. The supernatant was removed and subsequent steps were repeated for a second wash. The Qiagen DNeasy Blood & Tissue Kit protocol was then followed for DNA isolation.

#### Preparation of frozen organoids for DNA isolation

To prepare the cryopreserved organoid cultures for DNA isolation, the cryopreserved vial was thawed in a 37°C water bath for 1-2 minutes. The cell suspension was placed in a 5 mL tube with 3 mL of cold human wash medium with 0.1% BSA and centrifuged at 400 RCF for 5 min at 4°C. The supernatant was aspirated and the pellet was re-suspended in 200 µL ice-cold PBS with 0.1% BSA. The suspension was transferred to a 1.5 mL Eppendorf tube and triturated vigorously to blend organoids into solution. Ice-cold PBS with 0.1% BSA (1 mL) was added and the tube was centrifuged at 6000 RCF for 3 minutes at 4°C. The resulting supernatant was discarded and the Qiagen DNeasy Blood & Tissue Kit protocol was then followed for DNA isolation.

#### DNA fingerprinting

To assess sample quality and integrity and establish sample identity, 34-42 ng (15 ng/μL) of isolated DNA from each sample was aliquoted and used to generate a DNA fingerprint with the 96-SNP Advanta™ SampleID Genotyping Panel (Fluidigm, Inc). Briefly, the DNA samples were pre-amplified using the Advanta specific target amplification (STA) primers according to the manufacturer’s protocol and the amplified DNA along with the Advanta SampleID allele-specific (ASP) and locus-specific (LSP) primers were transferred to a Fluidigm 96.96 Dynamic Array Integrated Fluidics Circuit (IFC) for Genotyping. The IFC was processed on the Fluidigm Juno IFC Controller, and real-time PCR was performed on the Fluidigm Biomark HD System and the data were reviewed in the Fluidigm Genotyping Analysis Software.

### Histology and Immunohistochemistry

To determine whether morphological features are preserved in the organoids, we performed hematoxylin and eosin (H&E) and immunohistochemical (IHC) staining for MUC5AC (mouse monoclonal antibody, Ventana, Inc), the cancer epithelial marker cytokeratin-19 (CK19) (mouse monoclonal antibody, Ventana, Inc), and Ki-67 (rabbit monoclonal antibody, Ventana, Inc) in both the organoids and their parental tumor tissue. Organoids were revived from frozen organoid culture or generated from fresh tissue. Briefly, 500 μL of Dispase (5U/mL) diluted 1:5 in HBSS was added per well and incubated for 10 min at 37°C, 5% CO_2_ incubator. Organoid suspension from one confluent or two semi-confluent wells were harvested and transferred to a 1.5 mL Eppendorf tube and centrifuged at 1,500 rpm for 6 min.

The pellet was washed once with 1x PBS and resuspended in 1mL of 4% PFA and incubated at room temperature for 30 min or overnight at 4°C. The suspension was centrifuged as above and the pellet was washed twice with 1x PBS. To form a plug, the pellet was resuspended in 100 μL of warmed Histogel and allowed to polymerize at 4°C for 10 min. The organoid plug was removed from the Eppendorf tube and transferred to a 5 mL tube containing 2 mL of 70% ethanol and sent to the histology lab. Organoids were paraffin embedded and subjected to serial sections for H&E and IHC following per standard histology lab operating procedures.

#### Targeted DNA sequencing

As proof-of-concept, we evaluated the genomic similarity between a set of 4 tumor organoids and their primary parental tumor and normal tissue by isolating DNA and conducting targeted sequencing of 275 cancer-related genes using the QIAseq Human Comprehensive Cancer Panel. Forty nanograms of DNA was fragmented and used to construct whole-genome DNA libraries, followed by a single-primer extension-based target enrichment and library final amplification according to the manufacturer’s protocol. Following screening on the Agilent TapeStation 4200, the libraries were qPCR-quantitated using the Kapa Library Quantification Kit (Roche, Inc.), and the libraries were sequenced with a 2×150 base paired-end sequencing run on the Illumina NextSeq 500. After sequencing, the results were analyzed with the Qiagen QIAseq Targeted Sequencing Data Analysis Portal. Somatic mutations were determined by subtracting any variant observed in the normal sample from the passing variants detected in the parent IPMN or PDO samples. Identity was again confirmed by comparing genotypes at common polymorphisms.

## Results

Tissue samples from 15 unique IPMN surgical cases were processed to harvest organoids from paired IPMN tumor and adjacent normal pancreatic tissues. The cases included 9 male and 6 female patients ranging from 61 to 88 years of age at the time of resection, of which 40% were diagnosed due to incidental findings on computed tomography (CT) imaging (Figure 2A). The IPMN specimens were resected from varying locations in the pancreas including the pancreatic head, uncinate process, body, and tail. The IPMN cohort included 3 low-grade IPMNs, 2 moderate-grade IPMNs, 7 high-grade IPMNs, and 3 IPMNs associated with invasive carcinoma. Several distinct epithelial subtypes were characterized by histology and immunohistochemistry including pancreatobiliary, intestinal, and mixed subtypes (Figure 2B). Further details regarding select clinicopathological characteristics of the IPMN study cohort are listed in Table 1.

**Table 1.** Clinicopathological characteristics of the IPMN Study Cohort

|  | Total (n=15) | IPMN with<br>low-grade<br>dysplasia<br>(n=3) | IPMN with<br>moderate-grade<br>dysplasia<br>(n=2) | IPMN with<br>high-grade<br>dysplasia<br>(n=7) | Carcinoma<br>arising from<br>IPMN (n=3) |
| --- | --- | --- | --- | --- | --- |
| Median age (years) | 74 | 76 | 72 | 74 | 76 |
| Gender (%) |  |  |  |  |  |
| Male | 9 (60) | 2 (66.7) | 1 (50) | 3 (42.9) | 3 (100) |
| Female | 6 (40) | 1 (33.3) | 1 (50) | 4 (57.1) | 0 |
| Clinical presentation (%) |  |  |  |  |  |
| Symptomatic | 9 (60) | 2 (66.7) | 1 (50) | 4 (57.1) | 2 (66.7) |
| Incidental Finding | 6 (40) | 1 (33.3) | 1 (50) | 3 (42.9) | 1 (33.3) |
| Location (%) |  |  |  |  |  |
| Head | 7 (43.8) | 0 | 1 (50) | 4 (50) | 2 (66.7) |
| Uncinate process | 3 (18.8) | 1 (33.3) | 1 (50) | 1 (12.5) | 0 |
| Body | 3 (18.8) | 1 (33.3) | 0 | 2 (25) | 0 |
| Tail | 2 (12.5) | 1 (33.3) | 0 | 1 (12.5) | 0 |
| Diffuse | 1 (6.2) | 0 | 0 | 0 | 1 (33.3) |
| Biliary stent (%) |  |  |  |  |  |
| Yes | 1 (6.7) | 0 | 0 | 0 | 1 (33.3) |
| No | 14 (93.3) | 3 (100) | 2 (100) | 7 (100) | 2 (66.7) |
| Median CA 19-9 (U/mL) | 18.1 | 18.2 | 42.3 | 15.1 | 24.3 |
| Post-op tumor size (%) |  |  |  |  |  |
| <3 cm | 14 (87.5) | 2 (66.7) | 2 (100) | 7 (87.5) | 3 (100) |
| ≥3 cm | 2 (12.5) | 1 (33.3) | 0 | 1 (12.5) | 0 |
| Tumor stage (%) |  |  |  |  |  |
| T<sub>is</sub> | 5 (31.3) | 0 | 0 | 5 (62.5) | 0 |
| T1 | 2 (12.5) | 0 | 1 (50) | 0 | 1 (33.3) |
| T2 | 1 (6.2) | 0 | 0 | 0 | 1 (33.3) |
| T3 | 1 (6.2) | 0 | 0 | 0 | 1 (33.3) |
| Unspecified | 7 (43.8) | 3 (100) | 1 (50) | 3 (37.5) | 0 |
| Lymph node metastasis (%) |  |  |  |  |  |
| N0 | 9 (56.2) | 0 | 1 (50) | 5 (62.5) | 3 (100) |
| Unspecified | 7 (43.8) | 3 (100) | 1 (50) | 3 (37.5) | 0 |
| Duct involvement |  |  |  |  |  |
| Main duct | 4 (25) | 1 (33.3) | 1 (50) | 2 (25) | 0 |
| Branch ducts | 1 (6.2) | 0 | 0 | 1 (12.5) | 0 |
| Mixed | 3 (18.8) | 0 | 1 (50) | 2 (25) | 0 |
| None | 4 (25) | 1 (33.3) | 0 | 2 (25) | 1 (33.3) |
| Unspecified | 4 (25) | 1 (33.3) | 0 | 1 (12.5) | 2 (66.7) |
| Subtype |  |  |  |  |  |
| Pancreatobiliary | 6 (37.5) | 2 (66.7) | 1 (50) | 2 (25) | 1 (33.3) |
| Intestinal | 2 (12.5) | 0 | 0 | 1 (12.5) | 1 (33.3) |
| Foveolar/Intestinal | 1 (6.2) | 0 | 0 | 1 (12.5) | 0 |
| Intestinal/Pancreatobiliary | 1 (6.2) | 0 | 0 | 1 (12.5) | 0 |
| Intestinal/Gastric | 1 (6.2) | 0 | 0 | 1 (12.5) | 0 |
| Gastric/Pancreatobiliary | 1 (6.2) | 0 | 1 (50) | 0 | 0 |
| Unspecified | 4 (25) | 1 (33.3) | 0 | 2 (25) | 1 (33.3) |

**Figure 2.**
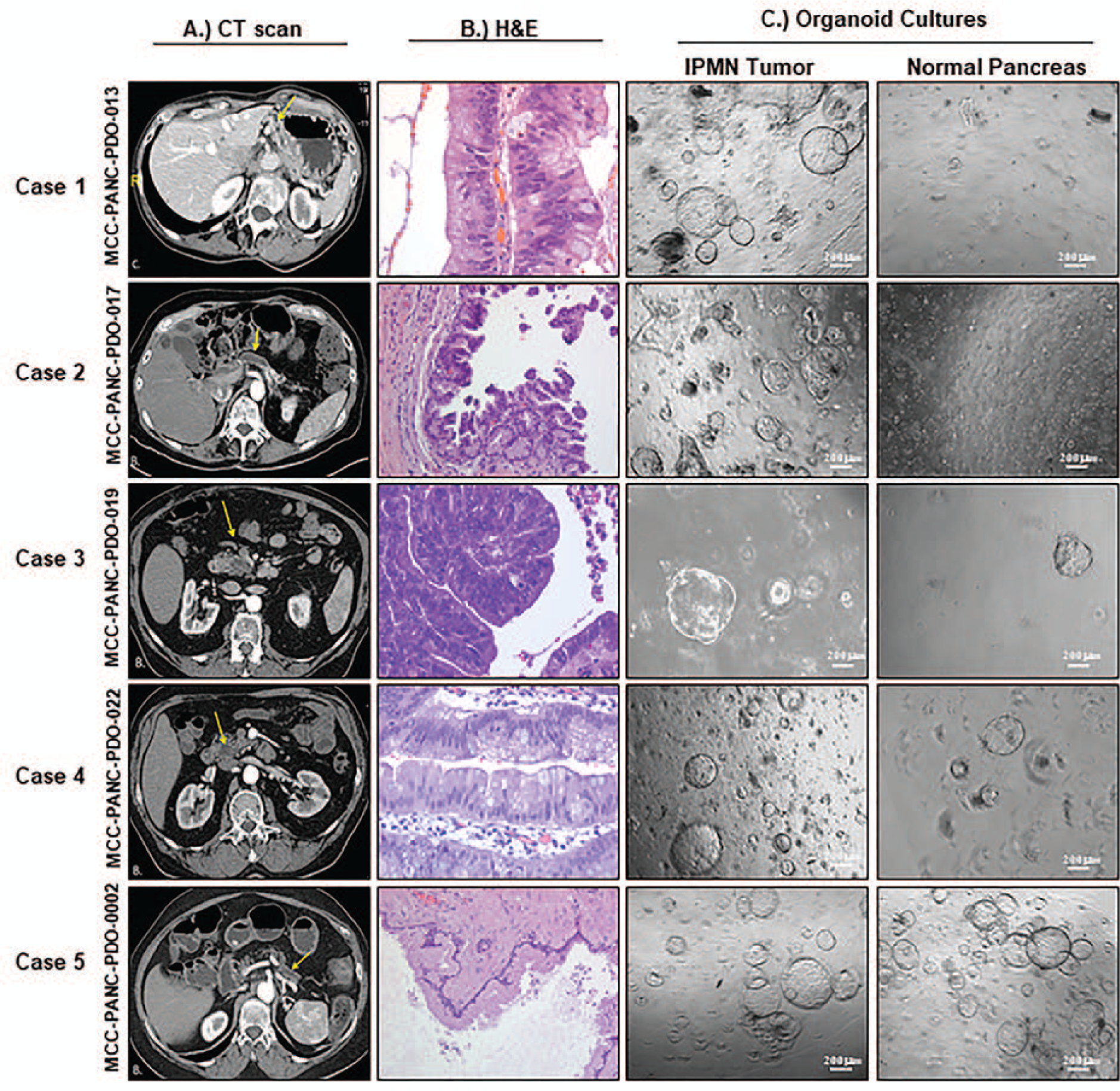
Generation of organoids from characterized human resected IPMN and normal tissues. A) Axial arterial phase preoperative computed tomography (CT) images of pancreatic IPMNs. B) Hematoxylin and eosin (H&E) images of resected IPMN tumor tissue. C) Bright-field micrographs (5x magnification) of organoid cultures derived from resected IPMN tumors and adjacent normal pancreatic tissue around day 7 or 8 unless noted otherwise. *Case 1 (MCC-PANC-PDO-013):* A. Moderate pancreatic ductal dilatation is present in the pancreatic tail (arrow). B. H&E (400X) shows high grade dysplasia of the mixed intestinal and gastric subtype characterized by pseudostratification with the nuclei remaining near the basal layer. The nuclei are vesicular with small nucleoli and visible goblet cells. *Case 2 (MCC-PANC-PDO-017):* A. Demonstrates diffuse irregular pancreatic main duct dilatation (arrows) compatible with main duct IPMN. B. H&E (200X) shows columnar cells with basally located nuclei that merge into neoplastic cells with pseudostratification and high-grade architectural features. *Case 3 (MCC-PANC-PDO-019):* A. Shows a hypodense pancreatic head mass with subtle internal arterial phase enhancement (arrow). B. H&E (400X) shows high grade dysplasia characterized by columnar neoplastic cells with eosinophilic cytoplasm. The nuclei overlap, are vesicular with nucleoli, and show loss of polarity and mitoses. C. Organoids were isolated from frozen tissue 33 days post-resection for this high-grade IPMN of pancreatobiliary subtype. *Case 4 (MCC-PANC-PDO-022):* A. Hypodense mass (arrow) with mild internal enhancement in the pancreatic head. B. H&E (400X) shows moderate grade dysplasia featuring papillary fronds lined by columnar, neoplastic cells, with pseudostratification and goblet cells. The nuclei are elongated and enlarged, with nuclear grooves. *Case 5 (MCC-PANC-PDO-0002):* A. Images through the pancreatic tail demonstrate diffuse dilatation of the main pancreatic duct (arrow). B. High power showing basally located nuclei. This field shows low grade dysplasia of pancreatobiliary subtype. C. Images taken at day 5. Scale bar, 200mm.

### Generation of IPMN organoid models

Among the 15 IPMN cases, two distinct IPMN tumor samples were resected from patient MCC-PANC-PDO-017 located at the pancreatic head and body, respectively, for a total of 16 tumor samples. Thirteen of sixteen IPMN tumor PDOs grew while thirteen of fifteen normal pancreatic organoids grew, for overall organoid generation success rates of 81% and 87%, respectively.

Cases with unsuccessful organoid development from IPMN tumor tissues were MCC-PANC-PDO-010, derived from an IPMN-pancreatobiliary subtype associated with invasive carcinoma; MCC-PANC-PDO-017, derived from an IPMN with high-grade dysplasia located at pancreatic body; and MCC-PANC-PDO-019, derived from an IPMN-pancreatobiliary subtype with high-grade dysplasia. Adjacent normal pancreatic tissues from MCC-PANC-PDO-017 and MCC-PANC-PDO-019 were also unsuccessful for organoid development.

Both normal pancreatic and IPMN organoids grew within 3 days. The majority of organoid cultures exhibited a cystic morphology (Figure 2C). Tumor cultures with other morphologic features included MCC-PANC-PDO-013 and MCC-PANC-PDO-017, which exhibited a mixed morphology characterized by both cystic and solid-filled shapes (Figure 2C). The organoid culture derived from frozen tissue, MCC-PANC-PDO-019, also exhibited a cystic morphology (Figure 2C).

We observed a different growth rate between the IPMN organoid cultures derived from fresh and frozen resected specimens. Organoids derived from fresh resected tissue grew at a faster rate than organoids derived from frozen resected tissue (Figure 3A). In terms of grade of differentiation, organoid cultures derived from resected high-grade IPMNs grew at a slower rate than organoid cultures derived from low and intermediate/moderate-grade IPMN specimens (Figure 3A).

**Figure 3.**
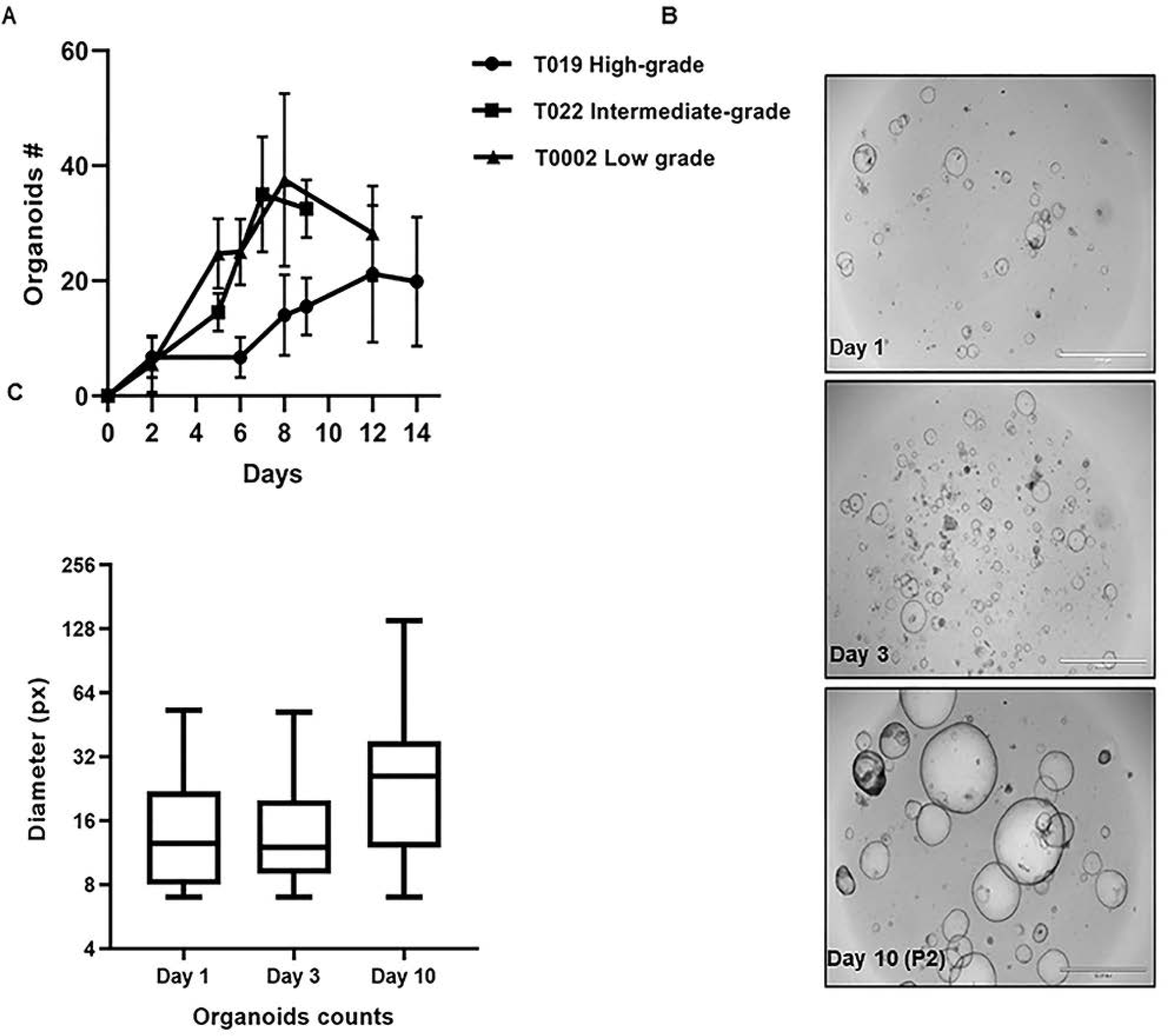
Expansion and biobanking of cryopreserved human IPMN organoids. Growth rate of IPMN Subtypes, Passaging and Reviving. (A) Growth curve of patient-derived organoid culture from MCC-PANC-T019 high-grade IPMN of pancreatobiliary subtype, isolated from cryopreserved tissue 33 days post resection; MCC-PANC-T022 intermediate-grade IPMN of pancreatobiliary subtype isolated from fresh resected tissue; and MCC-PANC-T0002 low-grade IPMN of pancreatobiliary subtype from fresh resected tissue (B) Patient-derived IPMN MCC-PANC-T005 organoid was revived 2 years post freezing, expanded, and passaged on Day 5 of culturing, then grown for an additional 10 days. Images were acquired on an EVOS FL Auto Imaging System, scale bar = 2mm (4x magnification). (C) MCC-PANC-T005 was analyzed by size (pxls) on the indicated days of culture.

### Establishment and Characterization of IPMN Organoids

We successfully established 15 tumor IPMN PDO out of 20 samples using a modified version of an established procedure for tissue digestion ^25^. This modified procedure uses wash media instead of feeding media for tissue digestion, making it a more cost-effective approach. These established IPMN PDO allowed long-term expansion and recovery after freezing. We were able to passage the IPMN PDO up to four times. Organoids from a second passage are shown in Figure 3B. After two passages PDO-T005 size (pxls) was analyzed using the EVOS FL Auto Imaging System (Figure 3C). We also successfully revived IPMN PDO that had been frozen for two years. Of note, we included Y-27632 Rho kinase inhibitor in the thawing media to help the organoid recover from the freeze/thaw stress; as others have described it as enhancing cells survival after cryopreservation^27 28^.

### Morphological Analysis of IPMN Tumor organoids and primary tumors

Comparison of 3D brightfield images, H&E, and IHC stains for MUC5AC, CK19, and Ki-67 for four IPMN parental tumor samples and the derived organoids reveal that morphological structures of the parental tumor tissue are generally preserved in the organoids (Figure 4 A-D). For example, an H&E of the parental tissue for a low grade IPMN corresponds to an organoid that shows clusters with a central core surrounded by neoplastic cells (Figure 4A) while parental tissue from a high grade IPMN reveals that the organoid grows in clusters (Figure 4B). Expression levels of MUC5AC and CK19 are similar in the parental tumor and organoid for several cases (Figures 4A-C) and remains independent of whether organoids were freshly generated, PDO-0246 (Figure 4C) or revived, PDO-002, PDO-005 and PDO-0002 (Figure 4D). The expression levels of Ki-67 staining is similar in both organoids and its parental tissue (Figure 4B-D).

**Figure 4.**
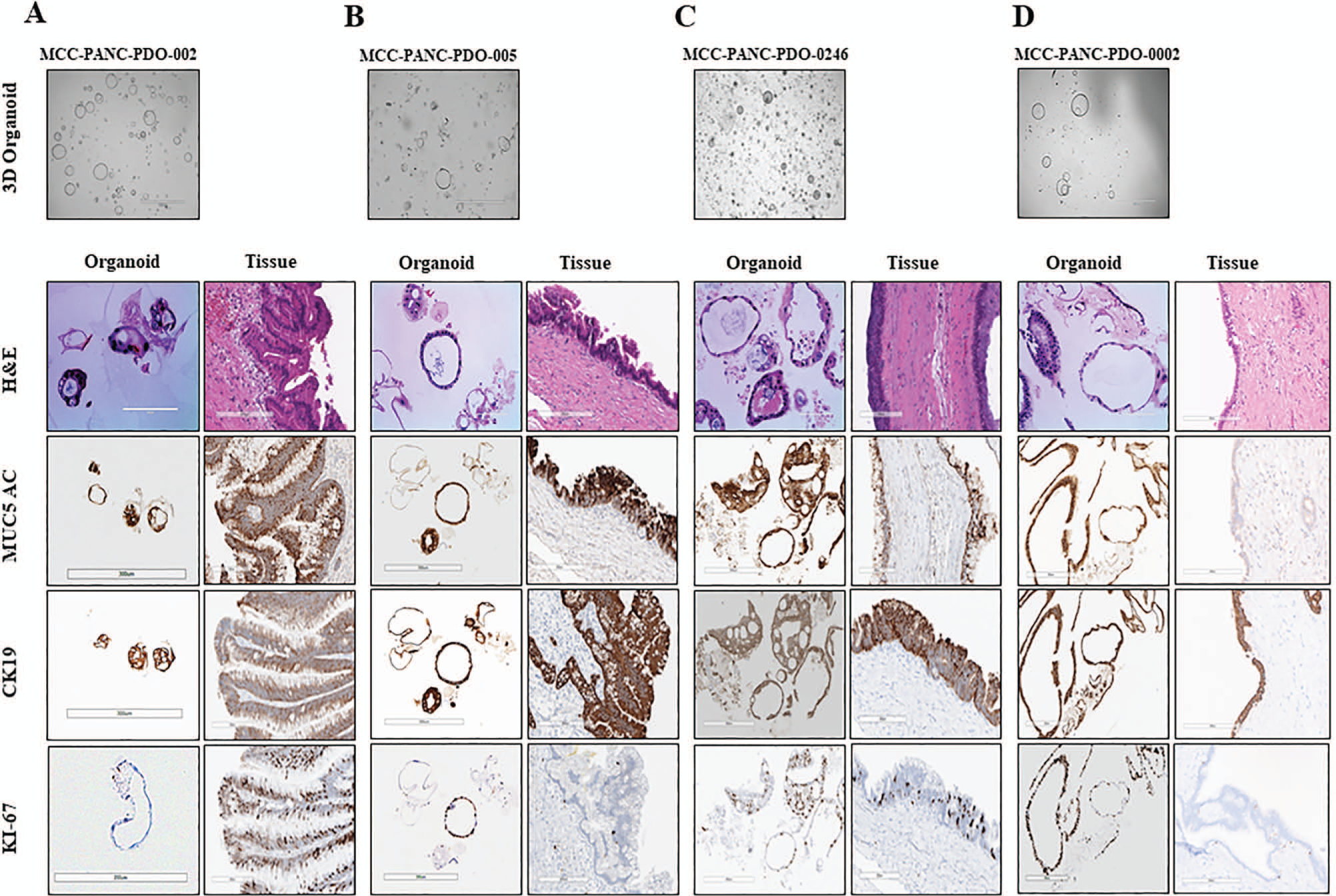
Morphological Analysis of IPMN Tumor Organoids and Primary Tumor. Morphological analysis of IPMN tumor organoids and primary tumors. Shown are 3D Brightfield images and H&E and immunohistochemical studies for MUC5AC, CK19 and Ki-67 for four IPMN tumor tissue samples and their derived organoids. A. MCC-PANC-PDO-002. The H&E of the tissue shows a low grade IPMN. The organoid shows clusters with a central core surrounded by neoplastic cells. Both the organoid and tissue strongly express MUC5AC, CK19. The tissue has a higher Ki-67 proliferation index than the organoid. B MCC-PANC-PDO-005. Tissue is a high grade IPMN. The organoid grows in clusters. The MUC5AC and CK 19 show a comparable expression level. The organoid cluster shows a higher Ki-67 than the tissue in this field. C. MCC-PANC-PDO-0246. The tissue shows low-grade IPMN. The MUC5AC, CK19 and ki-67 expression levels are similar in the organoid and tissue. D. MCC-PANC-PDO-0002. The tissue shows low grade IPMN with attenuated lining epithelium secondary to previous fine needle aspiration of the cyst. The CK19 expression levels are similar. The MUC5AC shows faint cytoplasmic expressions, but the expression in the organoid is strong. The organoid and tissue show similar expression for CK19. The Ki-67 is higher in the organoid sample.

### Organoid imaging & Automated counting

Images of the organoids were captured with a bright-field microscope at a magnification low enough to include the whole sample. The images were processed and analyzed via Image-Pro Premier software. Quantification of organoids can be challenging because they are grown in a 3D culture environment. Overlapping of the organoids makes it difficult for the available automated segmentation technique to give an accurate count. In order to develop a protocol for counting organoids in vitro, the Moffitt Imaging Response Assessment Team (IRAT) Core optimized the Circular Hough Transform (CHT) based algorithm^29^ to detect overlapping circular objects. We successfully quantitated organoids and segmented them by size (pxls) using this technique (Figure 5A-C).

**Figure 5.**
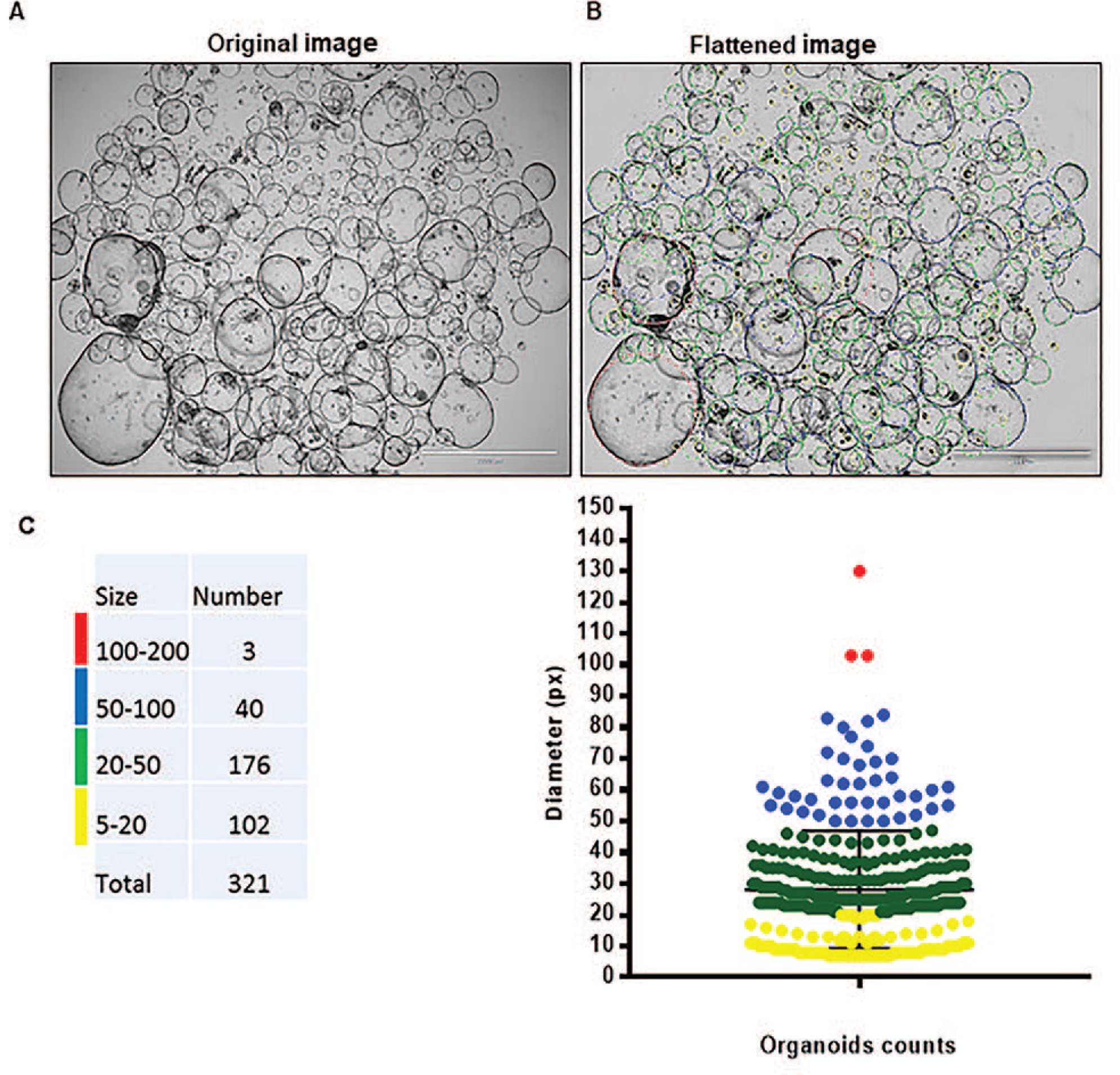
Automated counting of organoids using a Circular Hough Transform (CHT) based algorithm. Patient-derived organoid culture of low-grade IPMN of pancreatobiliary subtype (MCC-PANC-T0002) was revived, expanded, and passage twice. Image acquired at day 6 post passage. (A) Original image acquired on an EVOS FL Auto Imaging System (1.25x objective) (B) Flattened image and segmentation of detected objects. Scale bar = 2 mm. (C) Distribution of organoids based on count and size. Resulting organoid count color was coded by size (pxls).

### Genomic characterization of IPMN organoids

DNA was isolated from both normal and IPMN tumor organoids that showed successful growth, and was successfully performed on fresh and cryopreserved organoid samples. The concentration of DNA from normal samples ranged from 2.72 to 10 ng/µL and total DNA ranged from 44 to 223 ng. Their DIN scores ranged from 3.4 to 8. The concentration of DNA from IPMN tumor samples ranged from 2.32 to 33.2 ng/µL, the total DNA yields ranged from 131 to 672 ng, and the DIN scores ranged from 2.9 to 7.8.

All samples that underwent DNA fingerprinting passed quality control criteria and had single nucleotide polymorphism (SNP) genotype call rates that approached 100%. Additionally, the same identity for each PDO tumor-normal pair was verified, with nearly perfect correlations (r^2^>0.99) observed when comparing the allele intensity of samples in each pair. As expected, tumor and normal samples from unrelated individuals did not show correlated genotypes (r^2^<0.10).

In order to further characterize the IPMN organoids and determine whether their genomic profile matched their tissue of origin, 4 pairs of organoids and their corresponding paired parental tumor and normal tissue underwent targeted sequencing of cancer-associated genes using the QIAseq Human Comprehensive Cancel Panel. Average read depth of coverage ranged from 356x to 1201x, and average molecular barcode read depth of coverage was 20x to 170x. We observed an average of 41.5 somatic mutations in the targeted sequencing of organoids derived from IPMN tumors. Recurrent protein-altering somatic mutations were identified in key genes including *KRAS, GNAS, RNF43*, and *BRAF*, with somatic mutations identified in 2 (50%), 1 (25%), 1 (25%), and 1 (25%) of the 4 IPMN PDOs, respectively (Figure 6). Importantly, somatic mutations identified in the organoids were often found in the corresponding tumors (although tumor heterogeneity was observed), highlighting the potential for organoids to recapitulate genetic alterations commonly found in a pre-malignant disease state ^30,31^. For example, KRAS G12D and GNAS R844C were found in both the PDO and parental tumor for case 022, while *KRAS* mutations (G12D and G12V) observed in the PDO for case 0264 had evidence in the parental IPMN tumor, but at a lower frequency. For cases that did not have shared mutations between the parental tumor and PDO, it is possible that a different subclone was sequenced from the parent tumor tissue during the organoid culture process. An oncoprint showcasing protein-altering mutations in additional genes found in the PDO and parental tumors is featured in Supplementary Figure 1.

**Figure 6.**
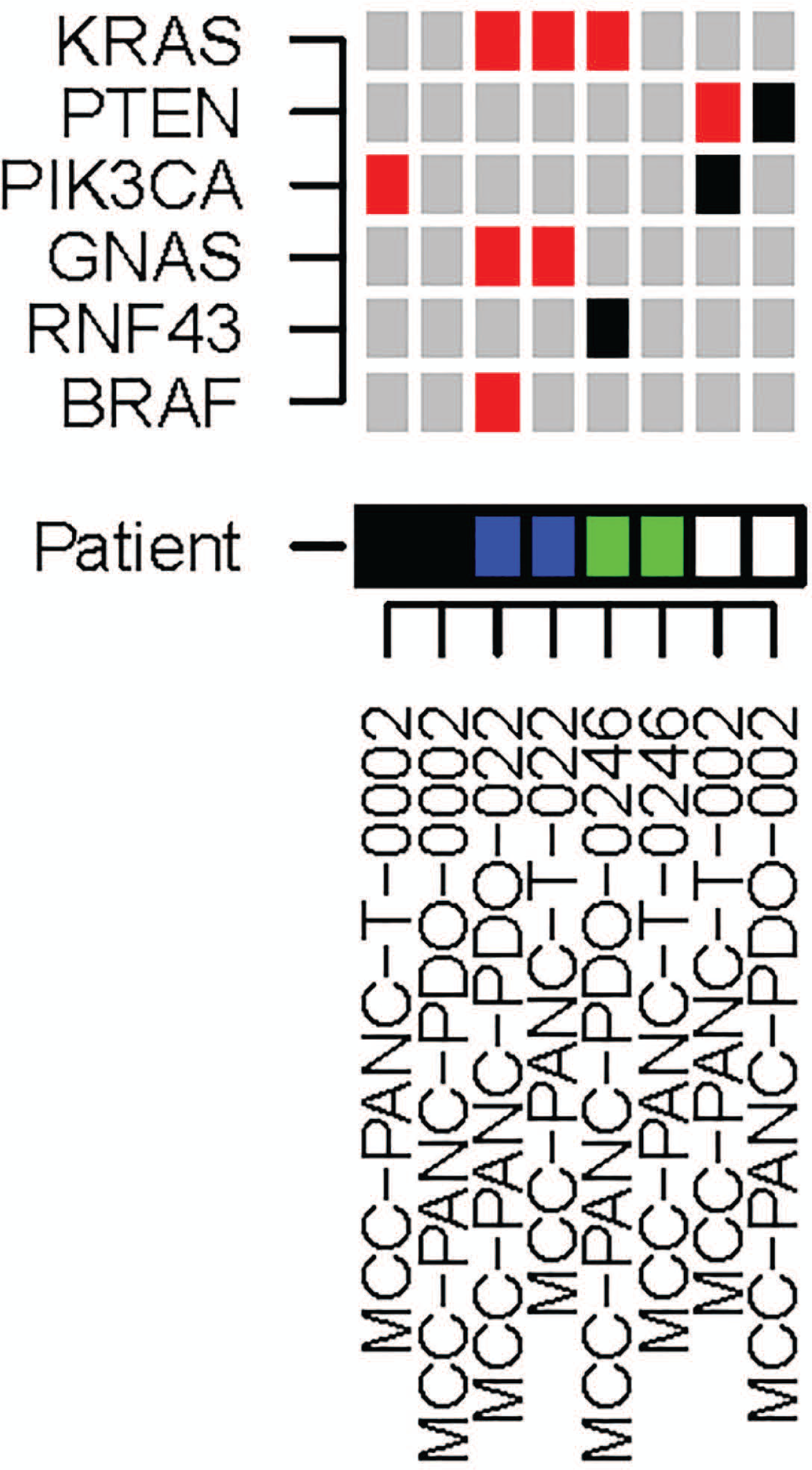
Characterization of IPMN tumor organoids by DNA sequencing. Oncoprint demonstrates that genes known to be mutated in IPMN tumors (*KRAS, PTEN, PIK3CA, GNAS, RNF43*, and *BRAF)* are mutated in a subset of paired IPMN parental tumor (T) and PDO samples from pathologically-confirmed low-grade (T002, T0002, T0246) and moderate-grade (T022) IPMN cases. Red boxes indicate protein-altering mutations, and black boxes indicate protein-truncating mutations.

## Discussion

We report the feasibility of generating viable patient-derived IPMN tumor and normal pancreatic organoids from newly resected and cryopreserved tissue at a success rate of 81% and 87%, respectively. Our outcomes are similar to previously-reported success rates of human pancreatic organoid models derived from normal and pancreatic ductal adenocarcinoma, which were up to 80% and 87% respectively^18, 22, 32-34^. The IPMN organoids were passaged, cryopreserved, and imaged with digital platforms^29, 35, 36^ which proved to be suitable to determine the number of overlapping organoids and organoid size over time. The IPMN PDO we generated exhibited a range of morphologies characterized by cystic and solid-filled spheres as previously described in organoid models derived from primary pancreatic cells and tissues^34, 37, 38^. Finally, histologic and genomic characterization of a subset of IPMN organoids and the paired parental tumor was successfully performed.

The establishment and expansion of pancreatic organoid cultures can require a two-week to five-month timeframe^34, 37, 38^, taking less time depending on the quantity of resected tissue available^39^. For example, over a 90% success rate has been reported for generation of colorectal cancer organoids with adequate amount of specimen^20^. In this investigation, IPMN organoid cultures were optimally established by a two-week incubation followed by passage and cryopreservation. Growth rate differences were observed between low-grade and high-grade IPMNs, with high-grade IPMNs growing at a slower pace than low-grade IPMNs. Although this may be linked to distinct genetic and stromal factors that may impede growth and may be specific to individual patients^39^, this observation was based on a small number of samples and warrants replication before drawing any firm conclusions.

We showed that the organoid culture model can recapitulate the morphological and mutational profiles of resected IPMNs, consistent with existing data on primary PDAC tumors^18-21^ and a recently published manuscript on IPMNs^23^. Discordance of somatic mutations was reported in several parental tumor-PDO pairs, suggesting a different subclone of parental tissue may have been sequenced compared to the subclone(s) that grew into organoid culture, consistent with prior studies which reveal driver gene heterogeneity in IPMNs^23, 40^.

There are several areas that need to be addressed before translational advances can be made. For example, it is important to assess factors that may have resulted in organoid isolation failure (including tissue growth requirements^41^ and slight deviations of protocols that may have impacted tissue processing after acquisition, tissue quality, and digestion time)^31^. Also, the Wnt ligands often do not possess sufficient activity to promote organoid growth; therefore, its monitoring is recommended to ensure optimal culture conditions^42^. The establishment of normal and tumor organoids under the same conditions may favor overgrowth of normal epithelial cells and halt organoid isolation^43^. High levels of genomic instability may lead to organoid apoptosis which may have posed a challenge upon generation and passage of organoid cultures^43^. Other biological factors such as sampling variation from heterogeneous tumors, evolution, natural selection due to culture conditions, stromal components, and tumor necrosis may have contributed to the characteristic mutational profile of the organoids^44^. Finally, despite advances in 3D culture techniques, long-term expansion remains a challenge ^22^. Difficulties reviving the organoids were encountered and lengthy optimization techniques were performed in order to revive and expand the IPMN organoids, making survival beyond 4 passages a challenge as organoids did not maintain their structure and viability. Of note, Huang et al ^23^ also reported survival for up to 4 passages in their new IPMN organoid living biobank. There is a possibility that IPMN PDO require additional manipulation of the culture condition that would favor their expansion using newly described modifications^31^.

Despite these challenges, the organoid platform may foster an improved understanding of the histopathology, genetic mechanisms, and stromal interactions of pancreatic cancer precursors and provide insight into novel chemoprevention and treatment approaches^45^. There is also potential to generate organoids from pancreatic biopsy specimens^46-48^ since fine needle biopsies can be a sufficient source of cells to isolate organoids within two weeks, with a success rate of 87%^33, 44^. The capacity to generate these preclinical models from surgical specimens and biopsies also has great potential to foster personalized medicine approaches to combine the high-throughput drug screening with the molecular assessment of organoid models in order to identify key diagnostic and therapeutic biomarkers that will predict which patients are at a higher risk of progression^49^. Furthermore, based on studies that have shown the ability of CRISPR/Cas9-mediated genome editing to modify driver genes in human pancreatic ductal organoids, opportunity exists to replicate the genetic mechanisms leading to pancreatic tumorigenesis^50-52^.

In summary, this is the largest reported IPMN PDO living biobank derived from human resected tissues that we are aware of. The generation, passage, cryopreservation, and characterization of these models demonstrates feasibility in this approach which may ultimately enable clinical implementation in favor of improving patient outcomes and chemoprevention studies aimed to halt progression of IPMN to invasive cancer.

## Acknowledgements

The research supported in this publication was supported in part by a Moffitt Cancer Center Team Science Award (awarded to JPB, DJ, GMD, MM, KJ, and D-TC), the James and Esther King Biomedical Research Program, Florida Department of Health (Grant #8JK02; awarded to JPB), and the Tissue Core, Molecular Genomics Core, Analytic Microscopy Core, the Imaging Response Assessment Team, and the Biostatistics and Bioinformatics Shared Resource at the H. Lee Moffitt Cancer Center & Research Institute, an NCI designated Comprehensive Cancer Center (P30-CA076292). The content is solely the responsibility of the authors and does not necessarily represent the official views of the sponsors or the H. Lee Moffitt Cancer Center & Research Institute.

Editorial assistance was provided by the Moffitt Cancer Center’s Scientific Editing Department by Dr. Paul Fletcher & Daley Drucker. No compensation was given beyond their regular salaries. The authors also wish to thank Warren Gloria, Karen Coley, Rithi Somesh Shivaram, Dennis Hall, and Vyacheslav Petrovskyy for their assistance with pathological review of tissues in the gross room.

