## Supplementary figures and images for "Establishing a Living Biobank of Patient-Derived Organoids of Intraductal Papillary Mucinous Neoplasms of the Pancreas"

### Supplemental Figure 1

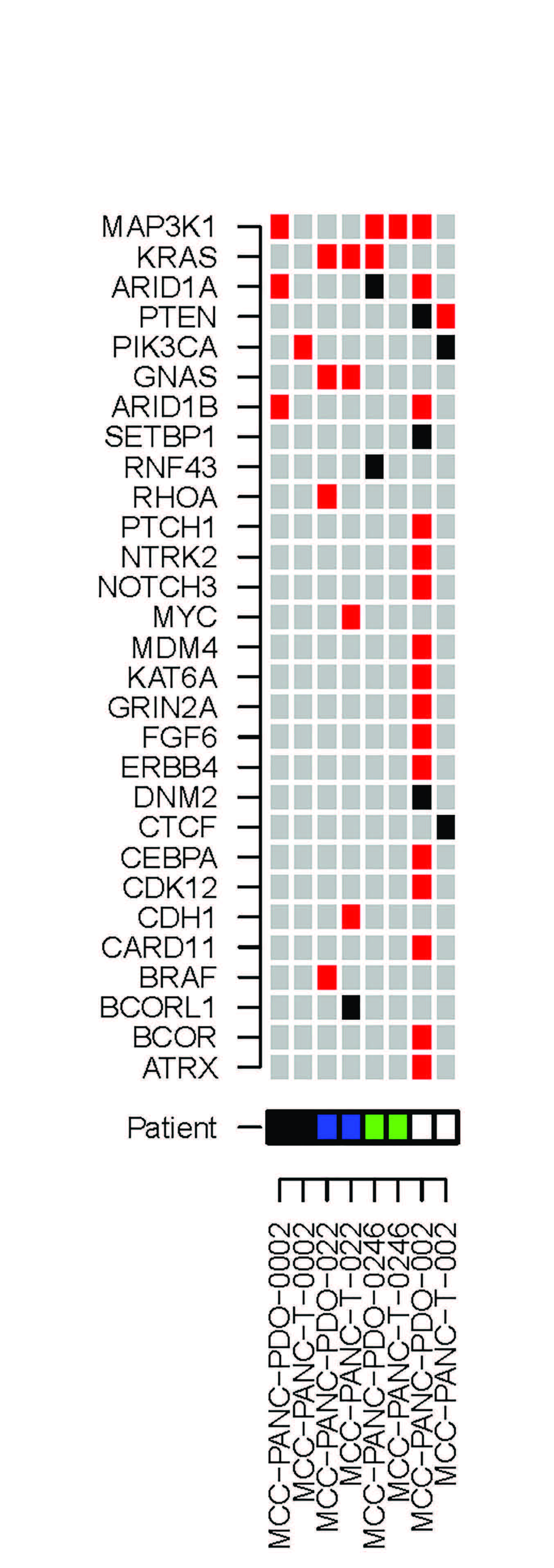
